# Fetal loss in pregnant rhesus macaques infected with high-dose African-lineage Zika virus

**DOI:** 10.1101/2022.03.21.485088

**Authors:** Lauren E. Raasch, Keisuke Yamamoto, Christina M. Newman, Jenna R. Rosinski, Phoenix M. Shepherd, Elaina Razo, Chelsea M. Crooks, Mason I. Bliss, Meghan E. Breitbach, Emily L. Sneed, Andrea M. Weiler, Xiankun Zeng, Kevin K. Noguchi, Terry K. Morgan, Nicole A. Fuhler, Ellie K. Bohm, Alexandra J. Alberts, Samantha J. Havlicek, Sabrina Kabakov, Ann M. Mitzey, Kathleen M. Antony, Karla K. Ausderau, Andres Mejia, Puja Basu, Heather A. Simmons, Jens C. Eickhoff, Matthew T. Aliota, Emma L. Mohr, Thomas C. Friedrich, Thaddeus G. Golos, David H. O’Connor, Dawn M. Dudley

## Abstract

Countermeasures against Zika virus (ZIKV), including vaccines, are frequently tested in nonhuman primates (NHP). Macaque models are important for understanding how ZIKV infections impact human pregnancy due to similarities in placental development. The lack of consistent adverse pregnancy outcomes in ZIKV-affected pregnancies poses a challenge in macaque studies where group sizes are often small (4-8 animals). Studies in small animal models suggest that African-lineage Zika viruses can cause more frequent and severe fetal outcomes. No adverse outcomes were observed in macaques inoculated with a low dose of African-lineage ZIKV at gestational day (GD) 45. Here, we inoculate eight pregnant rhesus macaques with a higher dose of African-lineage ZIKV at GD 45 to test the hypothesis that adverse pregnancy outcomes are dose-dependent. Three of eight pregnancies ended prematurely with fetal death. ZIKV was detected in both fetal and placental tissues from all cases of early fetal loss. Further refinements of this challenge system (e.g., varying the dose and timing of infection) could lead to an even more consistent, unambiguous fetal loss phenotype for assessing ZIKV countermeasures in pregnancy. These data demonstrate that high-dose inoculation with African-lineage ZIKV causes pregnancy loss in macaques and also suggest that ZIKV-induced first trimester pregnancy could be strain-specific.

**Author summary:** Although pregnant rhesus macaques are susceptible to infection with Zika virus (ZIKV), fetal phenotypes can be subtle and variable. Most macaque studies of ZIKV have involved infection with Asian-lineage viruses because these viruses caused Western Hemisphere outbreaks beginning in 2015. African-lineage ZIKV yields more severe adverse fetal outcomes in small animal models. Here, we provide evidence that pregnant macaques infected late in the first trimester using a high dose of African-lineage ZIKV have frequent ZIKV-associated pregnancy loss. This severe phenotype establishes a new model for evaluating countermeasures and reinforces the idea that African-lineage ZIKV infection may be a frequent cause of pregnancy loss in areas where it is endemic depending on the amount of transmitted virus during infection.

## Introduction

Zika virus (ZIKV) was first isolated in 1947 from the Zika forest in Uganda and was sporadically detected in Africa and Asia exclusively until two 21st-Century outbreaks: a 2007 outbreak in Micronesia and a 2013 outbreak in French Polynesia (1–3). Eventually, ZIKV spread to the Americas, where a dramatic increase in cases of microcephaly (fetal head size smaller than two standard deviations below average) coincided with a ZIKV outbreak in Brazil in 2015 (4–8). ZIKV gained global attention when scientists found many developmental abnormalities and neuropathological impacts causally associated with ZIKV infection *in utero*. Collectively, these are referred to as ‘congenital Zika syndrome’ (CZS).

There are two genetic lineages of ZIKV, Asian and African, that differ by approximately 12% at the nucleotide level and 3% at the amino acid level (9). Because ZIKV was neglected prior to the 2015 outbreak, most research has utilized Asian-lineage viruses that are representative of the outbreak in the Americas (2,10–12). The use of multiple doses, strains, and routes to infect macaques at different stages of pregnancy make it difficult to compare studies directly. Nonetheless, Asian-lineage viruses from Puerto Rico (ZIKV-PR), Brazil (ZIKV-BR), and French Polynesia (ZIKV-FP) can all cause adverse fetal outcomes, ranging from subtle to severe, with spontaneous fetal death occurring sporadically. In a meta-analysis combining data from several studies in different research centers, 28.6% of macaques infected with ZIKV-PR experienced pregnancy loss (13).

While this heterogeneity of outcomes is useful in dissecting the mechanisms of CZS, it is problematic for evaluating medical countermeasures such as vaccines and antivirals. Unreasonably large numbers of animals are needed to power experiments in which prevention of fetal loss is a primary outcome. The supply of macaques for research is limited, and this is especially acute for Indian-origin rhesus macaques which are the most widely captive-bred macaques in the United States. Several factors that restrict the use of pregnant Indian-origin rhesus macaques in research include export restrictions, low fecundity, and a small fertility window of four to five months per year (14). Furthermore, macaques used for ZIKV studies must be naive for other arthropod-borne flaviviruses and facilities often have limited infectious disease housing, a requirement for ZIKV-infected macaques.

Consequently, a pregnant macaque model where ZIKV causes consistent and severe adverse fetal outcomes to test therapeutics or prophylaxis would be desirable, enabling appropriately powered experiments with smaller group sizes. Mice infected with African-lineage ZIKV have increased fetotoxicity compared to mice infected with the Asian-lineage strains. For example, Ifnar1^−/−^ C57BL/6 mice infected with low-passage (five passages) African-lineage ZIKV strain ZIKV/Aedes africanus/SEN/ DAK-AR 41524/1984 (ZIKV-DAK) had more frequent fetal loss (100% vs. 53.2%) than mice infected with ZIKV-PR (15). Additionally, African-lineage strains of ZIKV showed an increase in peak viremia in both plasma and the brain that correlated with a decline in survival rates in adult immunocompromised mice (16,17). This increased pathogenicity has been recapitulated in studies that used a low-passage, African-lineage strain of ZIKV (ZIKV-DAK) and includes placental tissue damage *in vitro* (18) and more virus present in fetal organs in a porcine model (19). This also raises the worrisome possibility that if these results extend to primates, gestational African-lineage ZIKV could be a “silent” cause of pregnancy loss. In addition to posing a threat to human health, pregnancy loss could be especially consequential to endangered Great Apes that live in areas where ZIKV is endemic.

To determine if the high rate of early fetal loss associated with ZIKV-DAK in mice translates to primates, Crooks et al. inoculated four animals subcutaneously (SQ) with a low dose (1×10^4^ PFU) of ZIKV-DAK during the first trimester (gestational day 45). Given that first trimester ZIKV infections are more frequently associated with adverse fetal outcomes and ZIKV was detected in the MFI of the four low-dose ZIKV-DAK animals, it is surprising that all four animals had pregnancies proceed normally to study endpoint (20). Two other n=1 studies using Asian-lineage ZIKV (ZIKV-PR, ZIKV-CAM) have reported more frequent adverse pregnancy outcomes when macaques are infected SQ with approximately 10,000 times more virus than typically transmitted by mosquito bite (21–23). This provides some evidence that risk of fetal injury may be proportional to the amount of virus inoculated. While a dose that reflects infection by mosquito bite (1×10^4^-1×10^6^ PFU) is important to study natural pathophysiology of ZIKV infection, it is important to establish a model with higher reproducibility of significant adverse fetal outcomes to study countermeasures against the virus. Therefore, to further test the hypothesis that a high-dose of virus would increase fetal loss rate, we inoculated eight pregnant rhesus macaques SQ with 1×10^8^ PFU ZIKV-DAK. Here we observed pregnancy loss in three of eight pregnancies, establishing a macaque model for severe fetal outcomes following high-dose inoculation with African-lineage ZIKV.

## Results

### Pregnancy outcomes and maternal ZIKV infections

Eight ZIKV-naive rhesus macaque dams (046-101, 046-102, 046-103, 046-104, 046-105, 046-106, 046-107, 046-108) were inoculated subcutaneously (SQ) with a high dose, 1×10^8^ PFU/ml, of Zika virus/A.africanus-tc/Senegal/1984/DAKAR 41524 (ZIKV-DAK; GenBank: KX601166) at approximately GD 45 (range = GD 41-48) (Fig 1A and S1 Table). Data from these animals were compared to historical data collected from four dams infected with a low dose (1×10^4^ PFU/ml) of ZIKV-DAK and five dams mock-inoculated with sterile phosphate-buffered saline (PBS) (24). Physical exams of infected dams showed no ZIKV infection-associated symptoms such as rash or fever (S2A Fig). All animals had steadily increasing weights throughout pregnancy with the exception of 046-107 who lost 10.6% of her body weight between zero and 10 DPI (S2B Fig).

**Fig 1.**
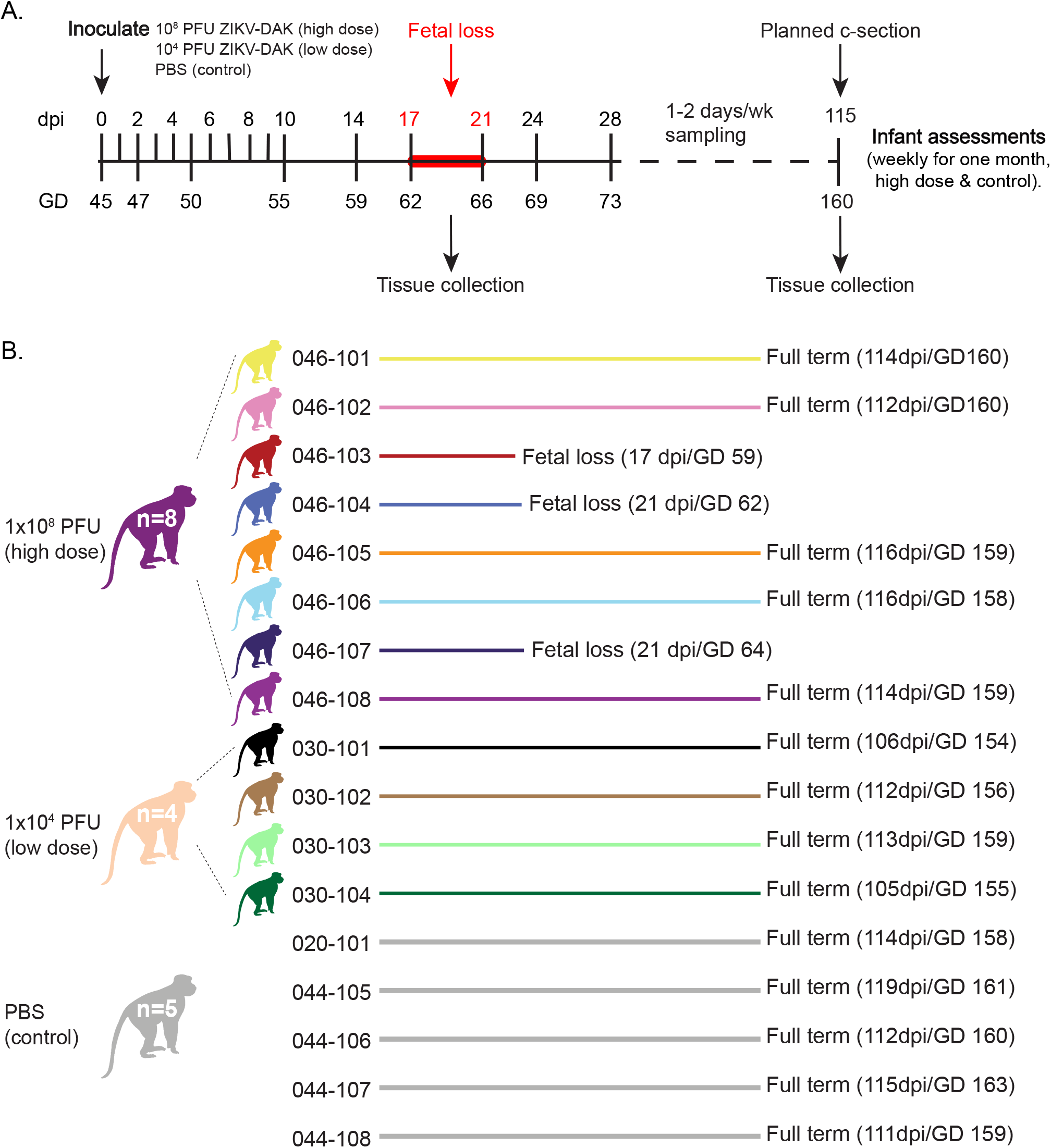
Experimental timeline and pregnancy outcomes. (A) Study timeline showing timing of infection and sampling. Samples were collected at each time point shown on the line. (B) Color representation and pregnancy outcomes of female rhesus macaques that were inoculated SC with either 1×10^8^ PFU ZIKV-DAK, 1×10^4^ PFU ZIKV-DAK, or PBS between GD 41 and GD 50. Mock animals were not assigned individual colors. Colors are used throughout the manuscript when describing the results from these animals and cohorts. Note that fetuses or infants will be color-matched to the appropriate dams.

ZIKV vRNA was detected in the plasma of all ZIKV-inoculated dams regardless of infection dose (Fig 2, S3 Fig). Peak plasma vRNA loads ranged from 9×10^2^-7×10^5^ copies/ml and were detected for all high dose dams between one and two DPI (Fig 2A, Fig 2C, Fig 2E, S3A Fig). Interestingly, two high dose dams (046-102 and 046-105) displayed qualitatively lower peak vRNA loads (approximately 1×10^3^ copies/ml) relative to the other dams that received a high dose of inoculum (S3A Fig). Viral RNA was also detected in the urine of two dams (046-101 and 046-104) and in the saliva of three dams (046-101, 046-107, and 046-108) during acute infection (S4 Fig). Dams inoculated with 1×10^4^ PFU had peak plasma vRNA loads that occurred later (between four and five DPI) and ranged from 1.33×10^5^-1.24×10^6^ copies/ml (Fig 2A, Fig 2C, Fig 2E, S3B Fig). Comparison of total plasma virus replication between high-dose and low-dose dams by t-test on the area under the curve (AUC) showed no significant difference between groups (t = −1.16; df = 3.11; p = 0.33) (Fig 2B). Similarly, comparison of peak plasma vRNA loads by t-test (t = 0.99; df = 3.91; p = 0.38) and plasma viral load duration by Wilcoxon ranksum tests with continuity correction (W = 6; p = 0.10) also showed no significant differences between high-dose and low-dose dams (Fig 2C, Fig 2D). A linear mixed effects model used to examine the time to peak plasma vRNA by group showed that dams that received a low dose of ZIKV-DAK had plasma vRNA loads that peaked significantly later than dams that received a high dose of the same virus (estimate = 2.50; std.error = 0.29; df = 10; t = 8.61; p < 2.0×10^−16^) (Fig 2E).

**Fig 2.**
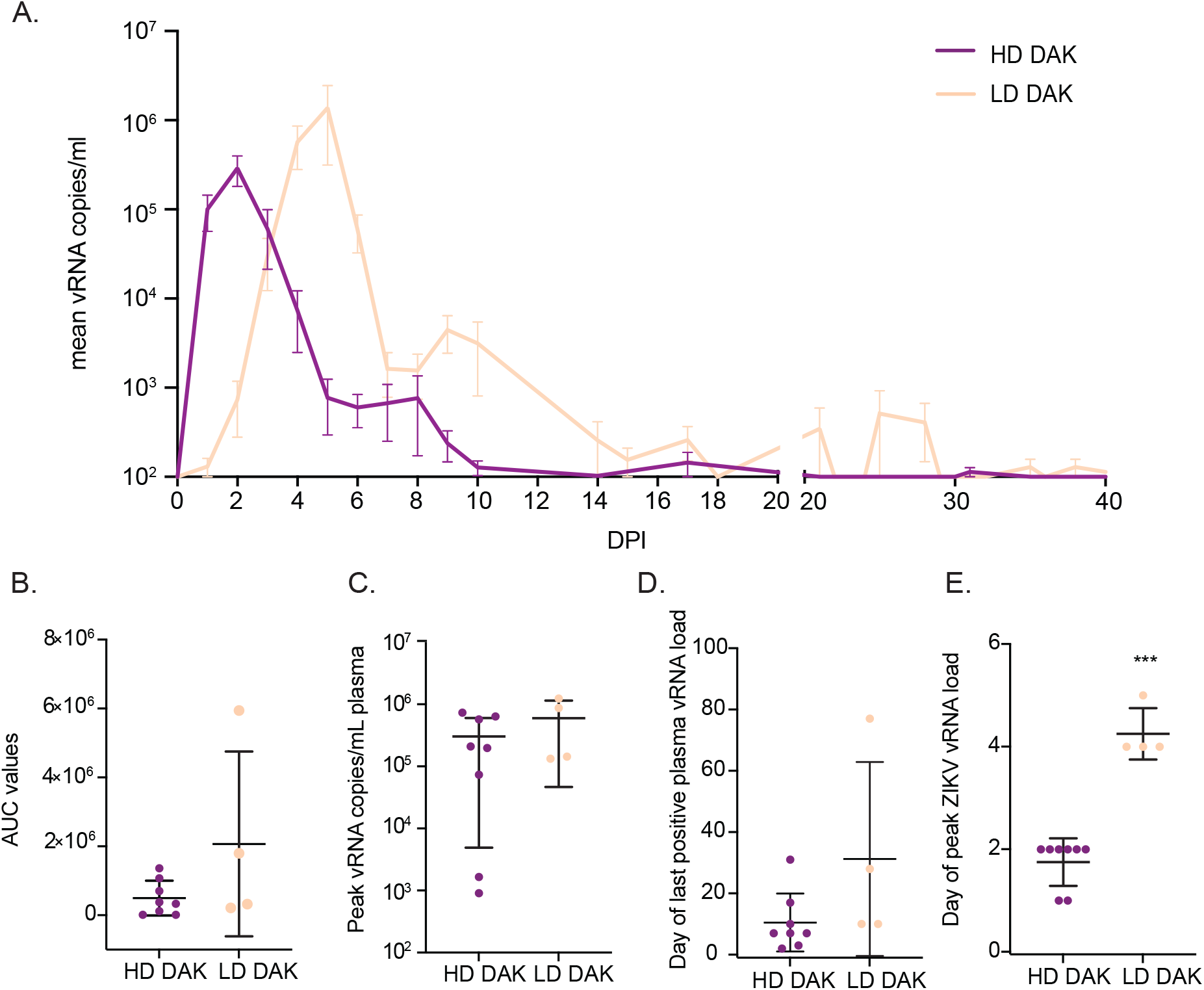
Replication kinetics of ZIKV-DAK in dams receiving a high dose (1×10^8^ PFU) or a low dose (1×10^4^ PFU) of ZIKV. (A) Mean viral loads through 40 DPI measured in plasma samples by ZIKV-specific RT-qPCR. Error bars represent the standard error of the mean (+/−SEM). (B) Comparison of plasma vRNA load area under the curve (AUC) for high-dose and low-dose inoculation groups. (C) Comparison of peak plasma vRNA load in copies/ml plasma. (D) Comparison of days post-infection (DPI) of the last positive plasma vRNA load. (E) Comparison of day post-infection (DPI) peak plasma vRNA load occurred. For all graphs in parts B-E, the mean value is represented by the wider black bar with error bars representing standard deviation (+/− SD). *** Represents a p-value of <0.001, while no asterisks represent no statistical difference at the 5% level.

Three of the eight high-dose dams had early fetal loss identified by ultrasound between GD 59 and 64 (Fig 1). One of the five dams that carried to near full-term lost her infant at five days of life due to postnatal complications not directly associated with ZIKV infection (failure to thrive). Because fetal demise occurred in three of eight high-dose pregnancies, we also explored whether maternal vRNA kinetics impacted pregnancy outcome by comparing overall maternal plasma vRNA load using AUC, peak plasma vRNA load, and duration between high-dose dams with early fetal loss and high-dose dams with fetuses that survived. We observed no significant differences between maternal plasma vRNA kinetics for any of these measures when high-dose animals were grouped by pregnancy outcomes (demise vs. survival) (S5 Fig).

### Maternal antibody response

Serum samples collected from all ZIKV-infected dams at zero, 21, and 28 DPI were assessed for neutralization capacity by 90% plaque reduction neutralization tests (PRNT_90_) on Vero cells. Results indicate that all eight high-dose animals and all four low-dose animals developed robust neutralizing antibody (nAb) responses following ZIKV-DAK infection (S6 Fig). Overall, PRNT_90_ titers at 21 or 28 DPI in dams with identified pregnancy loss and dams with viable pregnancies were not significantly different. ZIKV-specific IgM levels were measured in maternal serum samples in the high-dose ZIKV-DAK cohort by ELISA at zero, seven, 14, 17, and 21 or 24 DPI (S7 Fig). IgM levels peaked on days 14 or 17 post-infection for all eight dams. After 17 DPI, IgM levels began to decrease for all dams except 046-106.

### In-utero and at-birth measurements and observations

#### In-utero

Comprehensive ultrasounds were performed weekly to monitor growth, fetal heart rate, and observe changes in the fetus and placenta throughout pregnancy. As noted above, fetal death was determined in three of eight pregnancies by absent fetal heartbeat between 59 and 64 GD (046-503, 046-504, and 046-507) (S8 Fig). These three fetuses were found by ultrasound to have thick edematous skin which was attributed to fetal death in utero and subsequent autolysis (S9 Table). Other than possible early hydrops in one of the three fetuses, there were no abnormalities noted by ultrasound in the fetuses preceding demise. Fetal growth measurements of head circumference (HC), biparietal diameter (BPD), abdominal circumference, HC-femur length ratios, and BPD-femur length ratios, were also taken weekly or biweekly during routine ultrasounds. Neither ZIKV-DAK experimental group showed significant differences when compared to fetal measurements from mock-inoculated fetuses (S10 Fig, S11 Table, S12 Table).

#### At-birth

Measurements were taken at the time of birth for each infant. HC (p=0.5437), BPD (p=0.2506), and body weight (p=0.0849) of live born infants from high-dose dams were not significantly different when compared to infants from mock-inoculated dams (S13 Table). APGAR scores taken at one, five, and 10 minutes after birth were also not significantly different between infants from high-dose dams and mock-inoculated dams (S14 Table). Demographic characteristics at the time of birth such as gestational age, dam weight, dam age, and gender were compared between groups, and there were no significant differences (S15 Table).

### MFI tissue and maternal tissue vRNA loads

ZIKV RNA was detected by RT-qPCR in maternal/fetal interface (MFI) tissues from five of eight dams that received a high-dose infection with ZIKV-DAK and in all four dams receiving low-dose ZIKV-DAK (Fig 3). Two of the three dams with undetectable virus in the MFI (046-102, 046-105) also had reduced plasma viremia (Fig 3, S3A Fig). All three dams with early fetal loss, plus two others, had ZIKV RNA in the MFI. Viral RNA was found in all three layers of the placenta (decidua, chorionic villi, chorionic plate). Placentas from dams with early fetal loss had consistently higher viral loads than placentas from dams with viable pregnancies, and viral burden was similar across all three layers (Fig 3). Although ZIKV RNA is broadly detected in all three layers of the placenta, it is not uniformly detected throughout each cotyledon. Other MFI tissues analyzed by RT-qPCR included fetal membranes and interplacental collateral vessels. Although fetal membranes were positive for dams with and without fetal losses, more virus was detected in cases of fetal loss (Fig 3).

**Fig 3.**
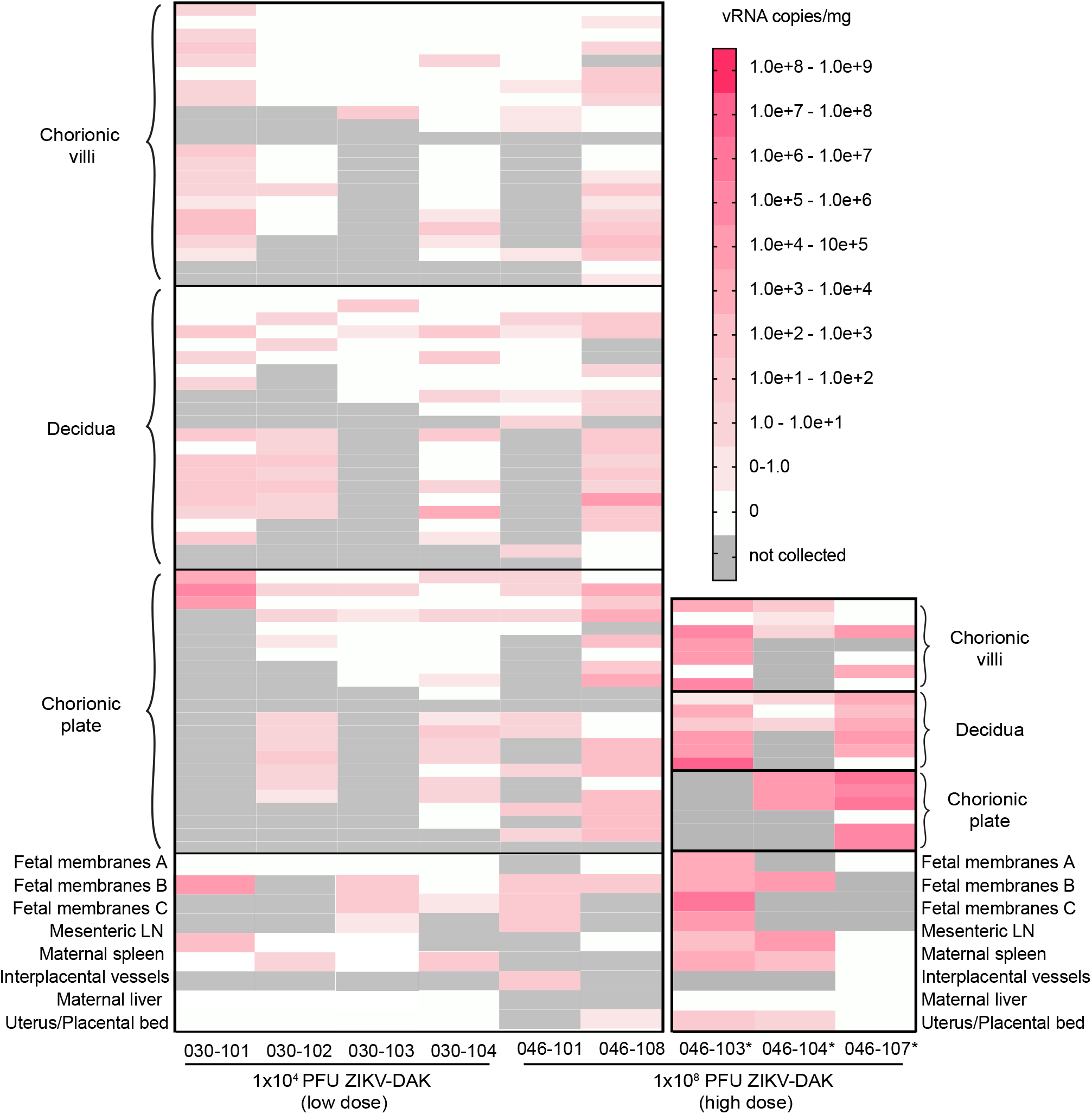
ZIKV RNA levels in maternal/fetal interface (MFI) tissues from high-dose and low-dose dams. Viral RNA was detected by ZIKV-specific RT-qPCR. Only animals with detectable virus in at least one tissue are shown on the plot. Dams with early fetal loss are marked with an asterisk.

Maternal tissues tested by RT-qPCR included mesenteric lymph node (LN), spleen, liver, and uterus at the placental bed (Fig 3). Mesenteric LN, maternal spleen, and the uterus/placental bed were all positive for ZIKV RNA in two of three dams with early fetal loss (046-103, 046-104). The only maternal tissues collected to test positive (10^0^-10^3^ copies/mg tissue) from dams with viable pregnancies were the uterus/placental bed (046-108), the mesenteric lymph node (030-101), and the spleen (030-102 and 030-104).

### Fetal viral loads

Viral RNA was detected by RT-qPCR in the fluids and/or tissues of all three fetuses following in utero death (Fig 4). Due to the early gestational time frame and small fetal size when demise occurred, organ and tissue samples were limited. Brain and eye from fetus 046-503 had the highest tissue vRNA burden observed. However, the highest overall vRNA loads were observed for the amniotic fluid and cerebrospinal fluid of fetus 046-507 (>1.00×10^7^ copies/mL) (Fig 4). Together, these results strongly suggest that vertical transmission occurred in all three instances of early fetal loss.

**Fig 4.**
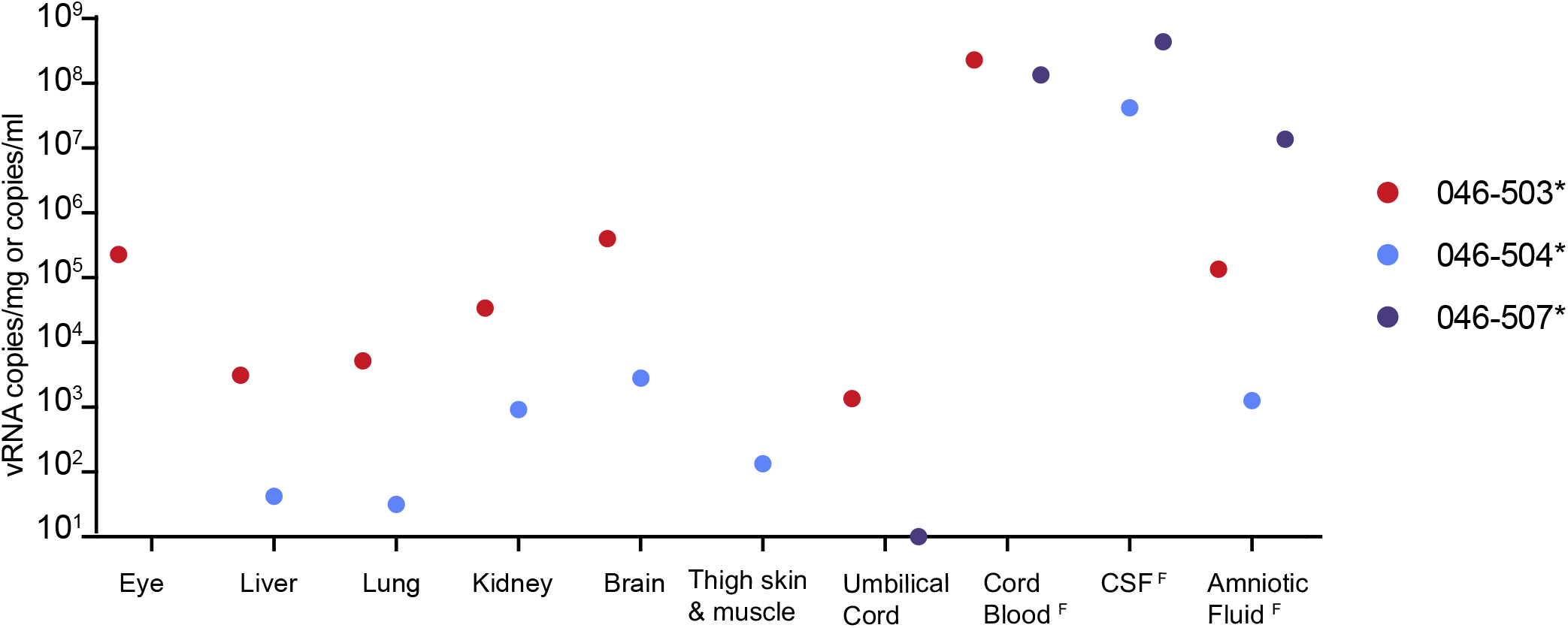
ZIKV RNA in fetal tissues and fluids detected by RT-qPCR from early fetal demise cases. Tissue samples were not available from 046-507. Fluid samples are distinguished from tissue samples with a superscript F and are measured in vRNA copies/ml.

### Fetal and placental pathology

For each of the three fetuses that died prior to planned c-section, significant tissue autolysis was identified which indicated that fetal death occurred at some point within the prior week. Unfortunately, fetal tissue autolysis made it difficult to determine whether pathological changes were specifically related to ZIKV infection.

Center cuts of all 12 placentas and cross-sections of individual cotyledons recovered from ZIKV-DAK infected dams (both high- and low-dose) showed increased evidence of pathology when compared to the four placentas recovered from uninfected dams, such as: chronic plasmacytic deciduitis, transmural placental infarction, and chronic histiocytic intervillositis (CHIV) (Table 1, S16 Fig, S17 Table).

**Table 1.**
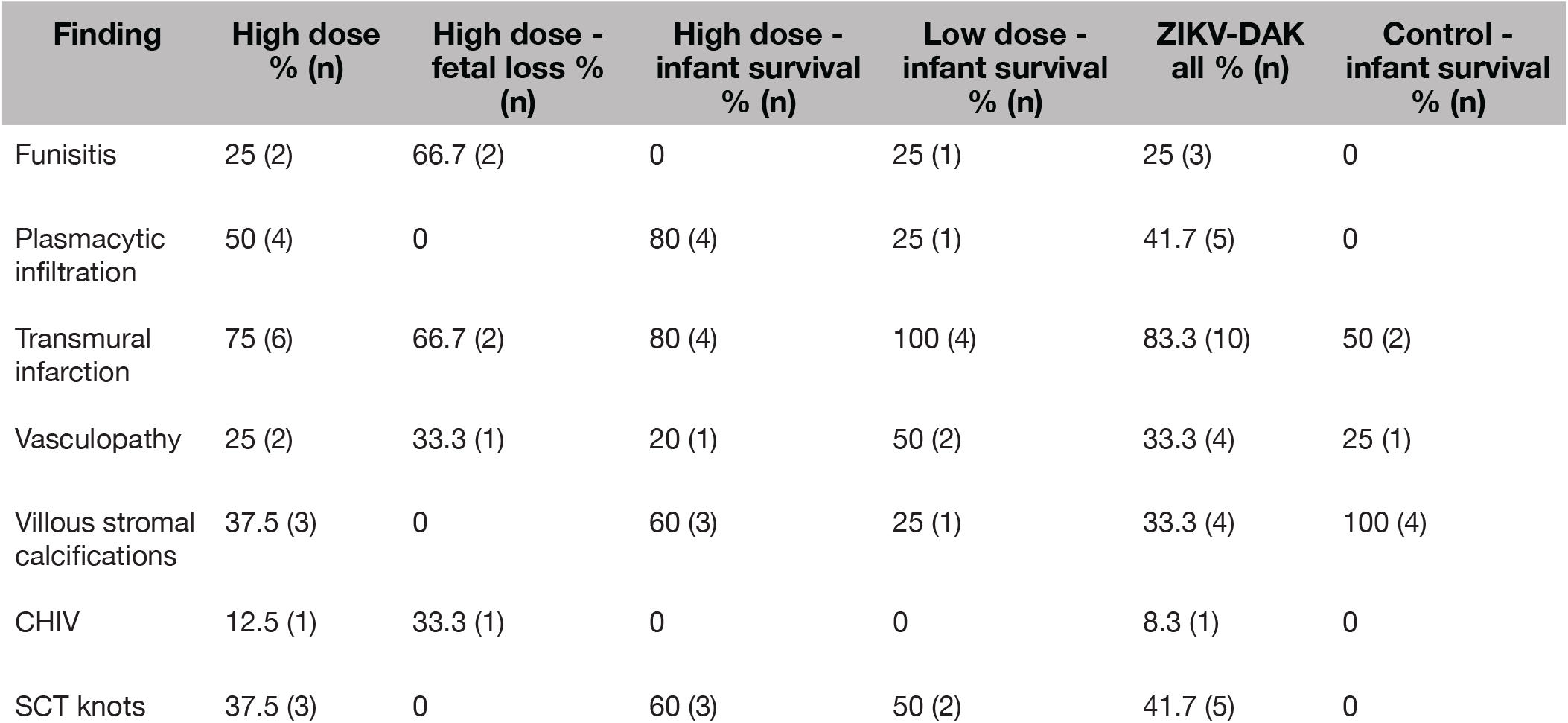
Histopathological analysis of maternal-fetal interface tissues. Animals are grouped in columns based on inoculation received and/or pregnancy outcome.

Overall, dams that received a high-dose infection with ZIKV-DAK had more pathological changes within the placenta than dams that received a low-dose infection with ZIKV-DAK, but comparisons to the low-dose animals were limited due to the smaller group size. Although both ZIKV-infected groups had more pathological changes identified in MFI tissues compared to mock-infected dams, the only finding unique to high-dose infection was CHIV identified in a case of fetal demise (046-107)(Table 1).

### Fetal and placental *in situ* hybridization (ISH)

To determine ZIKV RNA distribution in fetal and placental tissues from dams with early fetal loss (046-103, 046-104, 046-107), cross-sections from each placental cotyledon and multiple cross-sections from each fetus were analyzed by ZIKV RNA *in situ* hybridization (ISH). ISH revealed focal ZIKV infection in the placentas and fetuses in all three cases (Table 2)(Fig 5, Fig 6, Fig 7). Overall ZIKV detection was multifocal ZIKV RNA was detected throughout placental chorionic villi but restricted to the villous mesenchyme and absent from the syncytiotrophoblasts or basal plate (Fig 5). Multiple tissues of each fetus from fetal demise cases had vRNA detectable throughout the body. Tissues in all three fetuses with detectable vRNA included spinal cord and cerebral neuropil (Table 2, Fig 6, Fig 7, S18A-C, S18D-F). Viral RNA was also detected in the muscle, connective tissues, and periosteum (Table 2, Fig 6B, Fig 6C, S18J Fig, S18K Fig), muscle and mucosa of the intestines (Fig 6C, S18G Fig, S18I Fig), and the brainstem (Fig 7).

**Table 2.** Localization of ZIKV RNA throughout fetal tissues by ISH. Tissues and organs with positive ZIKV RNA are listed for each section. Tissues listed as ‘NC’ were not collected. Heart, lung, liver, and adrenal glands were either not collected or negative across all fetuses and are listed in this table.

| Tissue | 046-503 | 046-504 | 046-507 |
| --- | --- | --- | --- |
| Stomach | NC | NC | Positive |
| Intestines | Positive | Negative | Positive |
| Periosteum | Positive | Positive | Negative |
| Brainstem | Negative | Positive | Positive |
| Neuropil | Positive | Positive | Positive |
| Spinal cord | Positive | Positive | Positive |

**Fig 5.**
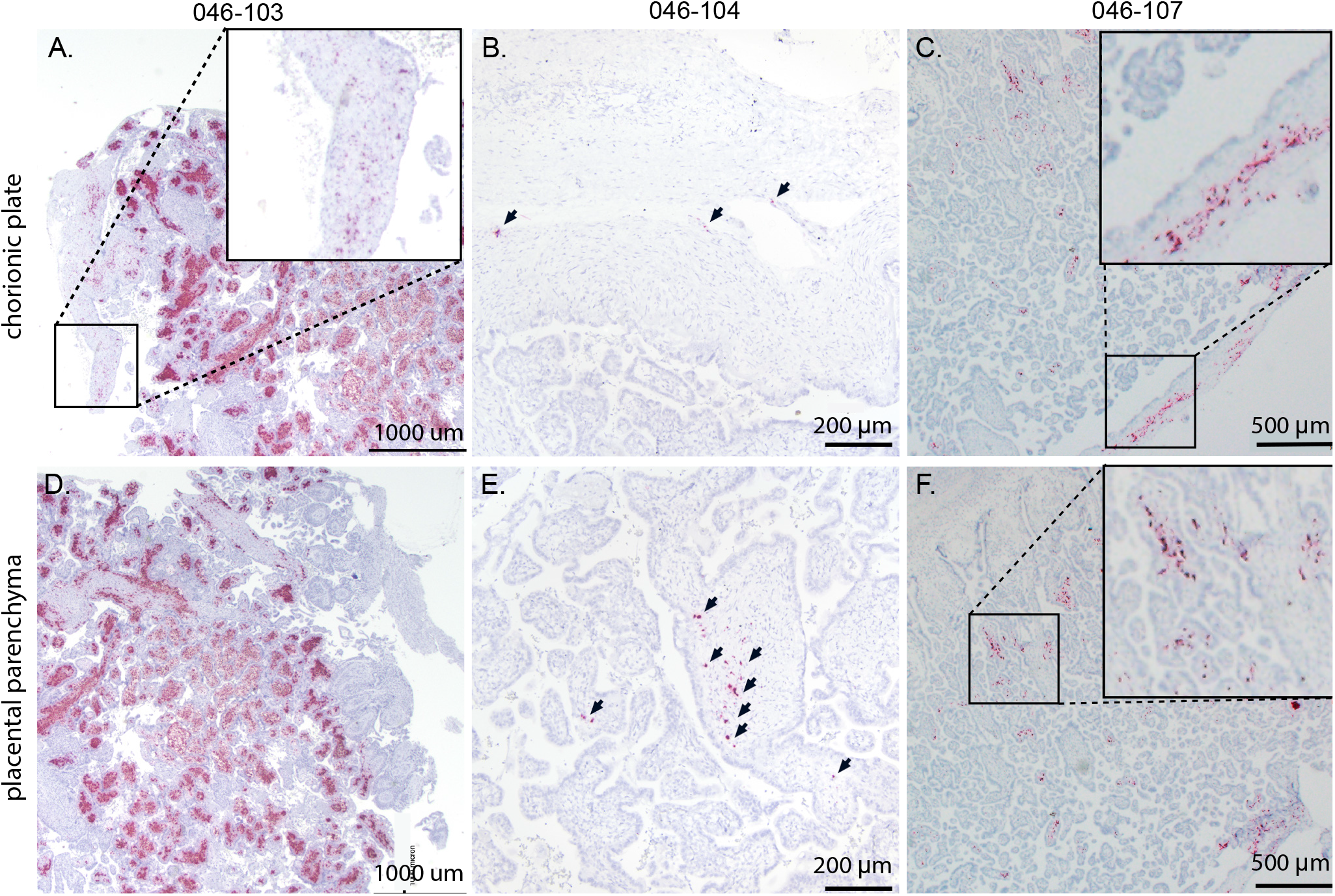
Detection of ZIKV RNA by in situ hybridization in MFI tissues of dams with early fetal loss. Representative images of ZIKV RNA distribution in MFI tissues from the three cases of early fetal loss. Numerous foci of ZIKV RNA were detected in the chorionic plate of (A) 046-103 (boxed), (B) 046-104 (arrows), and (C) 046-107 (boxed) as well as the placental parenchyma (villi) of (D) 046-103, (E) 046-104 (arrows), and 046-107 (boxed). ZIKV RNA is shown in red.

**Fig 6.**
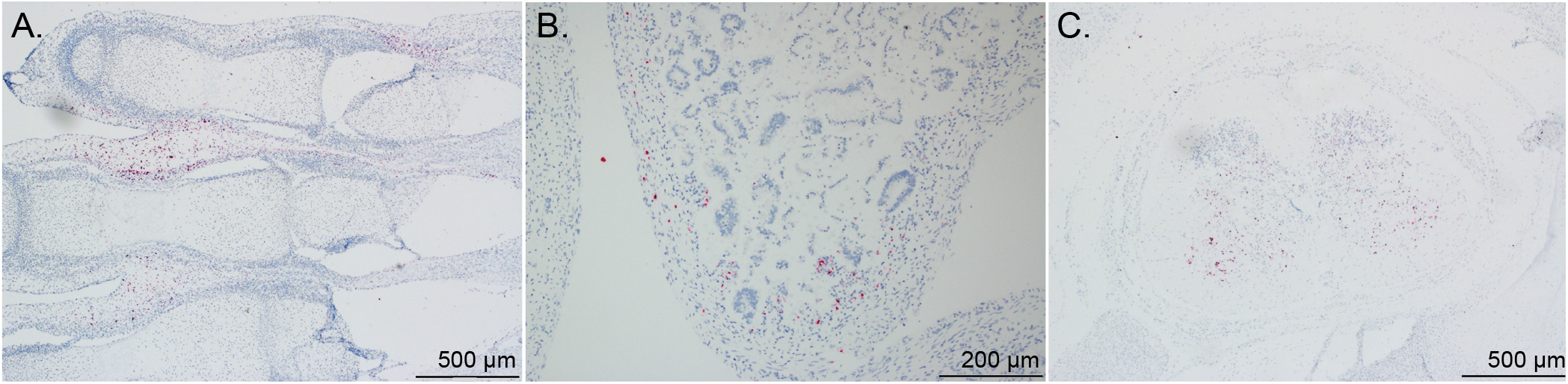
Detection of ZIKV RNA by in situ hybridization in fetal body tissues. Representative images of ZIKV RNA distribution in MFI and fetal tissues from the three cases of early fetal loss (excluding the head). Foci of ZIKV RNA were detected in (A) muscle and connective tissue of lower digits (046-503) and (B) of the intestines (046-507) as well as (C) the spinal cord (046-504). ZIKV RNA is shown in red.

**Fig 7.**
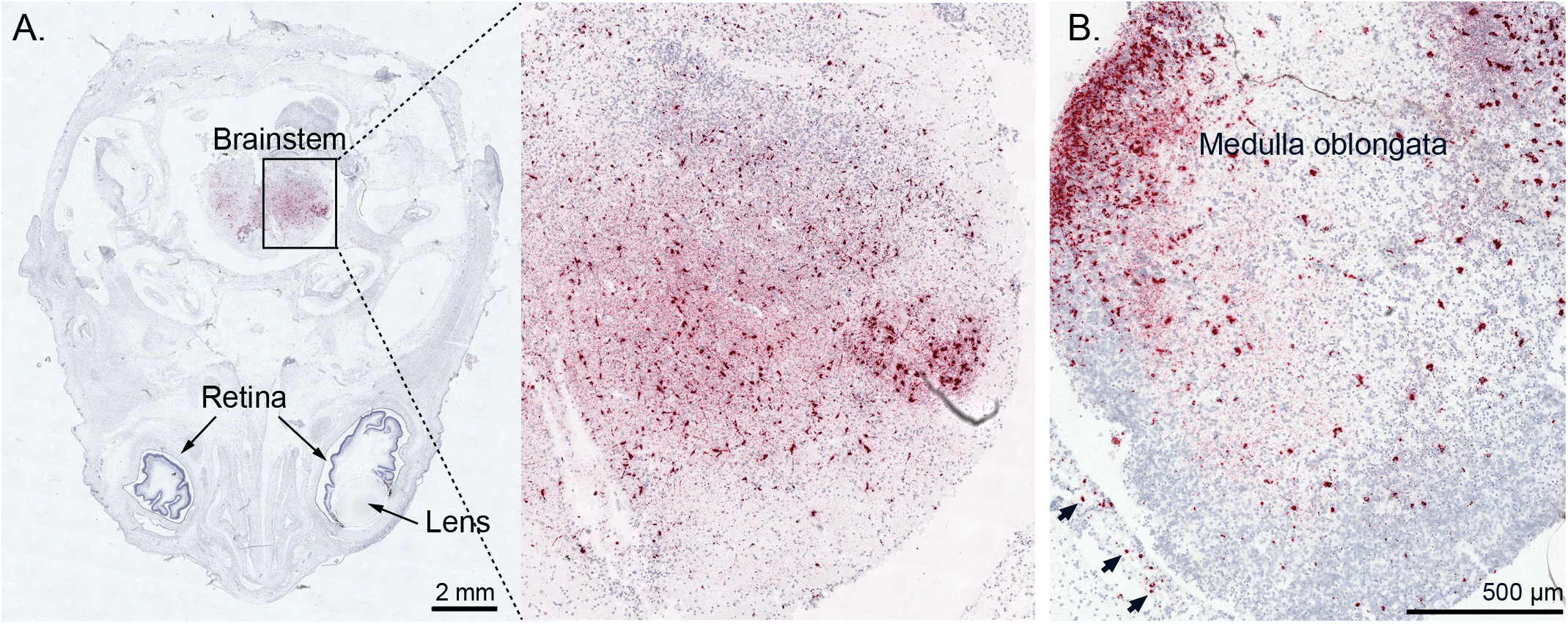
Detection of ZIKV RNA in fetal brainstem and head tissues from cases with fetal loss. ZIKV RNA was detected by in situ hybridization and is shown in red. (A) Foci of ZIKV RNA in the brainstem of fetus 046-504 and (B) foci of ZIKV RNA in the medulla oblongata region of the brainstem and the meninges (arrows) of fetus 046-507.

### Neonatal neurodevelopmental assessments and locomotion in the first month of life

Infants were monitored weekly during the first month of life using the Schneider Neonatal Assessment for Primates (SNAP) and Nodulus CatWalk. The four liveborn infants in the high-dose ZIKV-DAK group and the four infants in the control group scored similarly in the orientation construct at seven, 17, and 21 days of life (DOL), but at 28 DOL infants in the ZIKV-DAK group began to score worse (S19A Fig). The rate of motor development was also found to be slower for infants in the ZIKV-DAK group as compared to controls (S19B Fig). Although interesting, the differences in the orientation and motor development constructs are not significant (S21 Table). Assessment of motor coordination, balance, and gait patterns using the Noldus CatWalk revealed ZIKV-DAK infant walking patterns were more hetero-geneous than control infants at 14 and 21 DOL (S20C Fig, S22 Table). Specifically, ZIKV-DAK infants had a transitional gait pattern, or limb dragging (S20C Fig). This transitional, limb-dragging pattern, is distinct from the diagonal walking pattern seen in the Control group that is standard for developing infant macaques.

## Discussion

We found that increasing the dose of ZIKV-DAK increased the adverse fetal outcome rate to three in eight pregnancies, up from zero of four low-dose infections with ZIKV-DAK and two of 18 low-dose infections with ZIKV-PR. The primary adverse outcome we observed was fetal demise between 17 and 21 DPI. The consistent timing of fetal demise relative to ZIKV-DAK infection suggests a common cascade of pathologic events that culminate in fetal death. This consistent timing has two important implications. First, it creates a tractable system for evaluating therapeutics. Pregnancies that survive beyond this window are likely to survive to full term, or approximately 165 gestational days. So, less than half of a typical macaque pregnancy (45 GD inoculation plus approximately 30 days to assess fetal loss) is necessary to ascertain this outcome. Second, this rapid pregnancy loss, if extrapolated to higher-order primates including humans, would coincide with miscarriage in late first or early second trimester, a time when many pregnacies are spontaneously lost. This could explain why the impact of African-lineage ZIKV on pregnancies was not identified in the decades since the original discovery of this virus (25).

A key limitation to extrapolating these results to humans and other primates is that the subcutaneous (SQ) inoculation dose of 1×10^8^ PFU is higher than typically delivered by an infected mosquito, which is estimated to be 1×10^4^ to 1×10^6^ PFU (22,23,26). The unusually short time to peak viremia (approximately 2 days) observed following our high-dose SQ inoculations artificially shortens therapeutic window, which is typically 5 days for mosquito to macaque ZIKV transmission (26). However, use of therapeutics after 30 days can improve infection outcomes in unique cases of prolonged viremia (27). It should be noted that even if this model is not ideal for testing post-exposure therapeutics, it may still be valuable for unpacking mechanisms of fetal demise and understanding the neurologic impact of ZIKV-DAK on surviving infants with in utero exposure. It is also highly valuable for testing prophylaxis options.

The detailed examination of the three fetal losses could provide early clues to the mechanism(s) of fetal death. Detection of relatively high vRNA loads by RT-qPCR in fetal neural tissue and widespread presence of ZIKV RNA throughout the neural parenchyma (neuropil) by ISH in all three cases of early fetal loss suggest neurotropism for ZIKV-DAK (Fig 4, Fig 7, Table 2, S18A-C). Additionally, the presence of ZIKV RNA in the brainstem and spinal cord is consistent with human case studies that have identified spinal cord damage in cases of CZS with and without fetal loss, as well as brainstem hypoplasia (Fig 6D, Fig 7, Table 2, S18D-F Fig)(28–30). Given the evidence that ZIKV infiltrated the fetal CNS in all cases of early fetal loss, we anticipated that surviving ZIKV-DAK exposed macaque infants (046-501, 046-502, 046-505, 046-506) would display early neurodevelopmental deficits consistent with CZS. However, data collected from weekly neurodevelopmental assessments of surviving infants during the first month of life suggested no significant differences between groups (S19 Fig, S20 Fig, S21 Table, S22 Table). Future studies will need to explore emerging developmental trends beyond the first month of life to determine long-term impacts in infants prenatally exposed to ZIKV-DAK.

Collectively, these results suggest that infection with low-passage African-lineage ZIKV late in the first trimester is associated with fetal demise, but only when a higher dose of virus is administered. This particular model is useful for asking questions about the overall pathogenesis of infection and mechanisms of demise, as it yielded two different pregnancy outcomes across animals that received the same high-dose inoculations. Furthermore, because fetal loss does not occur in all cases using this model, we are able to begin to study the developmental impact of in-utero ZIKV-DAK exposure. Although an increased rate of adverse fetal outcome is useful for future intervention studies where cohort sizes are limited by constraints on pregnant rhesus macaques, a frequency of 37.5% may still not be sufficient to power studies of countermeasures adequately. Future intervention models must therefore build off of this model to further increase rates of adverse fetal outcomes, perhaps by infecting earlier in the first trimester when the fetus could be more vulnerable to viral infection.

## Materials and methods

### Study design

Eight rhesus macaques (*Macaca mulatta*) were identified, confirmed pregnant by ultrasound, and challenged during the first trimester at approximately GD 45 (term 165 ± 10 days) with 1×10^8^ PFU of an African-lineage ZIKV (ZIKV-DAK) administered subcutaneously (SQ). Macaque dams utilized in the study were free of Macacine herpesvirus 1, Simian Retrovirus Type D (SRV), Simian T-lymphotropic virus Type 1 (STLV), and Simian Immunodeficiency Virus as part of the Specific Pathogen Free (SPF) colony at WNPRC. Maternal health and pregnancies were monitored throughout the infection. Blood samples were collected for isolation of plasma and peripheral blood mononuclear cells (PBMC) from all dams prior to ZIKV challenge on days −4 and zero, daily post-challenge from days one to seven, then twice weekly until resolution of maternal plasma vRNA load, and once weekly until study completion (Fig 1A). Serum was collected on days −4 and zero pre-challenge, then on days two, four, seven, 10, 14, 24, 28, and 30 post-challenge, and then weekly thereafter. Urine was passively collected from the removable cage bottom pans below the animals’ cages at available time points. Saliva swabs were collected from dams on days zero to four and seven to 10 post-challenge, and then at all blood collection time points thereafter. Data collected from these eight dams were compared to data from dams previously infected with a low dose (1×10^4^ PFU) of ZIKV-DAK, as well as dams mock-inoculated with 1X phosphate-buffered saline (1X PBS) comparably sampled as controls (24).

### Care and use of macaques

The macaques used in this study were cared for by the staff at the Wisconsin National Primate Research Center (WNPRC) in accordance with recommendations of the Weatherall report and the principles described in the National Research Council’s Guide for the Care and Use of Laboratory Animals (31). The University of Wisconsin - Madison, College of Letters and Science and Vice Chancellor for Research and Graduate Education Centers Institutional Animal Care and Use Committee approved the nonhuman primate research covered under protocol number G006139. The University of Wisconsin - Madison Institutional Biosafety Committee approved this work under protocol number B00000117. All animals were housed in enclosures with required floor space and fed using a nutritional plan based on recommendations published by the National Research Council. Animals were fed a fixed formula, extruded dry diet with adequate carbohydrate, energy, fat, fiber, mineral, protein, and vitamin content. Macaque dry diets were supplemented with fruits, vegetables, and other edible objects (e.g., nuts, cereals, seed mixtures, yogurt, peanut butter, popcorn, marshmallows, etc.) to provide variety to the diet and to inspire species-specific behaviors such as foraging. To further promote psychological well-being, animals were provided with food enrichment, structural enrichment, and/or manipulanda. Environmental enrichment objects were selected to minimize chances of pathogen transmission from one animal to another and from animals to care staff. While on study, all animals were evaluated by trained animal care staff at least twice each day for signs of pain, distress, and illness by observing appetite, stool quality, activity level, and physical condition. Animals exhibiting abnormal presentation for any of these clinical parameters were provided appropriate care by attending veterinarians. Prior to initial viral infection and comprehensive ultrasounds, macaques were sedated using ketamine anesthesia and monitored regularly until fully recovered from anesthesia. Animals were not sedated for regular blood draws and fetal heart rate checks. When not sedated, animals were placed in a table-top restraint device to allow for sample collection.

### Viral Infection

Zika virus strain Zika virus/A.africanus-tc/Senegal/1984/DAKAR 41524 (ZIKV-DAK; GenBank: KX601166) was originally isolated from *Aedes africanus* mosquitoes in Senegal in 1984. One round of amplification on *Aedes pseudocutellaris* cells, followed by amplification on C6/36 cells, followed by two rounds of amplification on Vero cells, was performed by BEI Resources (Manassas, VA) to create the stock (five total passages) (15). Raw FASTQ reads (BioProject: PRJNA673500) and a FASTA consensus sequence (BioProject: PRJNA476611) of the challenge stock of ZIKV/Aedes africanus/SEN/DAK-AR-41524/1984 are available at the Sequence Read Archive. For virus inoculations, ZIKV-DAK stock was diluted to 1×10^8^ PFU in 1ml of 1X phosphate buffered saline (PBS) and delivered to each dam subcutaneously (SQ) over the cranial dorsum via a 1ml luer lock syringe.

### Ultrasonography and fetal monitoring

Fetal growth and viability were monitored every seven to ten days by ultrasound and doppler. Measurements such as heart rate (HR), biparietal diameter (BPD), head circumference (HC), femur length, and abdominal circumference (AC) were obtained. Fetal heart rates were monitored twice weekly throughout gestation to confirm viability. Mean growth measurements were plotted against mean growth measurements and standard deviations from rhesus macaques at specific gestational ages, collected by Tarantal et al. (32). To contextualize measurements collected before GD 50, the standard growth curve was extrapolated. Ultrasound interpretations were provided by a maternal-fetal medicine specialist.

### Temperature and body weight measurement

Rectal temperatures and body weights of the dams were collected throughout the study. WNPRC veterinary staff were consulted in determining whether elevation of an individual animal’s body temperature beyond reference ranges was clinically significant.

### ZIKV RNA isolation from plasma, urine, and saliva

Plasma and PBMCs were isolated from EDTA-treated whole blood by layering blood on top of ficoll in a 1:1 ratio and performing centrifugation at 1860 x rcf for 30 minutes with brake set at one. Plasma and PBMCS were extracted and transferred into separate sterile tubes. R10 medium was added to PBMCs before a second centrifugation of both tubes at 670 x rcf for eight minutes. Media was removed from PBMCs before treatment with 1X Ammonium-Chloride-Potassium (ACK) lysing buffer for five minutes to remove red blood cells. An equal amount of R10 medium was added to quench the reaction before another centrifugation at 670 x rcf for eight minutes. Supernatant was removed before freezing down of cells in CryoStor CS5 medium (BioLife Solutions) for long term storage in liquid nitrogen freezers. Serum was obtained from clot activator tubes by centrifugation at 670 x rcf for eight minutes or from serum separation tubes (SST) at 1400 x rcf for 15 minutes. Urine was passively collected from the bottom of animals’ housing, centrifuged for five minutes at 500 x rcf to pellet debris, and 270 ul was added into 30 ul DMSO followed by slow freezing. Saliva swabs were obtained and put into 500 ul viral transport media (VTM) consisting of tissue culture medium 199 supplemented with 0.5% FBS and 1% antibiotic/antimycotic. Tubes with swabs were vortexed and centrifuged at 500 x rcf for five minutes. Viral RNA (vRNA) was extracted from 300 uL plasma, 300 uL saliva+VTM, or 300 uL urine+DMSO using the Maxwell RSC Viral Total Nucleic Acid Purification Kit on the Maxwell 48 RSC instrument (Promega, Madison, WI).

### Maternal-fetal interface (MFI) collection and dissection from near full term pregnancies

Cesarean sections were performed to deliver full term infants. The placenta and associated membranes were harvested and immediately placed in sterile petri-dishes. Each placental disc was weighed and measured and then maintained on wet ice prior to same-day dissection. A full-thickness center-cut was collected from each placental disc and fixed with 4% PFA for histologic evaluation. The remainder of each placental disc was washed with 1X PBS and separated into cotyledons. A center-cut was taken from each cotyledon and placed into a biopsy cassette. Biopsy cassettes were fixed with 4% PFA for future analysis. Each cotyledon was divided into three layers: decidua, parenchyma, and chorionic plate. Two samples from each layer were collected and stored in either 750ul VTM or 1mL RNAlater. Tissues in VTM were frozen immediately after collection and stored at −80°C. Tissues in RNA later were left to sit in solution for 24 hours at 4°C, after which RNAlater was aspirated off and the tissues were stored at −80°C prior to vRNA isolation.

### Maternal-fetal interface (MFI) collection and dissection from first trimester fetal loss

Following early fetal demise a cesarean section was performed and the conceptus was immediately placed in a sterile petri dish. Fetal and maternal-fetal interface (MFI) tissues were evaluated by WNPRC veterinary pathologists and the tissues collected included full thickness center sections from each placental disc, three sections of amniotic/chorionic membrane from each placental disc, three sections of decidua from each placental disc, and one section of each of the following: fetal membranes, umbilical cord, maternal liver, maternal spleen, mesenteric lymph node (LN), uterus/placental bed, fetal liver, fetal lung, fetal kidney, fetal brain, fetal skin/muscle from thigh, fetal eye, fetal spleen, fetal upper limb, fetal chest, and fetal skull with brain. Two samples from each tissue section were also collected and stored in either 750ul VTM or 1mL RNAlater for vRNA assessment and future analysis. Tissues in VTM and RNAlater were stored respectively as described above.

### ZIKV RNA isolation from tissue samples

RNA was recovered from RNAlater-treated tissue samples using a modification of the method described by Hansen et al. (33). Briefly, up to 200 mg of tissue was disrupted in TRIzol Reagent (Thermo Fisher Scientific, Waltham, MA) with stainless steel beads (2×5 mm) using a TissueLyser (Qiagen, Germantown, MD) for three minutes at 25 r/s twice. Following homogenization, samples in TRIzol were separated using bromo-chloro-propane (Sigma, St. Louis, MO). The aqueous phase was collected into a new tube and glycogen was added as a carrier. The samples were washed in isopropanol and ethanol-precipitated overnight at −20°C. RNA was then fully re-suspended in 5 mM Tris pH 8.0.

### Viral RNA quantification by RT-qPCR

Viral RNA was quantified using a highly sensitive RT-qPCR assay based on the one developed by Lanciotti et al. (2008) (34), though the primers were modified to accommodate both Asian and African-lineage ZIKV. RNA was reverse transcribed and amplified using the TaqMan Fast Virus 1-Step Master Mix RT-qPCR kit (Invitrogen) on the LightCycler 480 or LC96 instrument (Roche, Indianapolis, IN) and quantified by interpolation onto a standard curve made up of serial tenfold dilutions of in vitro transcribed RNA. RNA for this standard curve was transcribed from a plasmid containing an 800 bp region of the ZIKV genome that is targeted by the RT-qPCR assay. The final reaction mixtures contained 150 ng random primers (Promega, Madison, WI), 600 nM each primer and 100 nM probe. Primer and probe sequences are as follows: forward primer: 5’- CGYTGCCCAACACAAGG-3’, reverse primer: 5′-CCACYAAYGTTCTTTTGCABACAT-3′ and probe: 5′-6-carboxyfluorescein-AGCCTACCTTGAYAAG-CARTCAGACACYCAA -BHQ1-3’. The reactions cycled with the following conditions: 50°C for five minutes, 95°C for 20 seconds followed by 50 cycles of 95°C for 15 seconds and 60°C for one minute. The limit of quantification of this assay is 100 copies/ml.

### Histology

Tissues collected for histology were fixed in 4% paraformaldehyde for one to four days depending on size and degree of autolysis prior to sectioning, processing, and embedding in paraffin. Paraffin sections were stained with hematoxylin and eosin (HE). Photomicrographs were taken using Olympus BX46 Bright field microscope (Olympus Inc.,Center Valley, PA) with an attached Spot flex 15.2 64Mp camera (Spot Imaging) and were captured using Spot Advance 5.6.3 software.

### IgM ELISA

Serum samples from zero, seven, 14, and 21 DPI were thawed at room temperature and added to an IgM fc-capture antibody conjugated plate (AbCam ab213327). HRP-conjugated ZIKV antigen (NS1) was added after the addition of serum, followed by addition of TMB and stop solution. The plate absorbance was read at dual wavelengths of 450nm and 600nm, IgM concentration is measured in AbCam units relative to the kit cut-off control (sample absorbance multiplied by 10, then divided by the absorbance of the cut-off control to yield AbCam Units).

### PRNT

Titers of ZIKV neutralizing antibodies (nAb) were determined for days zero, 21, and 28 post-infection using plaque reduction neutralization tests (PRNT) on Vero cells (ATCC #CCL-81) with a cutoff value of 90% (PRNT_90_) (35). Briefly, ZIKV-DAK was mixed with serial two-fold dilutions of serum for one hour at 37°C prior to being added to Vero cells. Neutralization curves were generated in GraphPad Prism (San Diego, CA) and the resulting data were analyzed by nonlinear regression to estimate the dilution of serum required to inhibit 90% Vero cell culture infection (35,36).

### *In situ* hybridization (ISH)

ISH probes against the ZIKV genome were commercially purchased (cat# 468361, Advanced Cell Diagnostics, Newark, CA). ISH was performed using the RNAscope® Red 2.5 kit (cat# 322350, Advanced Cell Diagnostics, Newark, CA) according to the manufacturer’s protocol. After deparaffinization with xylene, a series of ethanol washes, and peroxidase blocking, sections were heated with the antigen retrieval buffer and then digested by proteinase. Sections were then exposed to the ISH target probe and incubated at 40°C in a hybridization oven for two-hours. After rinsing, ISH signal was amplified using the provided pre-amplifier followed by the amplifier conjugated to alkaline phosphatase l, and developed with a Fast Red chromogenic substrate for 10 minutes at room temperature. Sections were then stained with hematoxylin, air-dried, and mounted prior to imaging using a Leica DM 4000B microscope equipped with a Leica DFC310FC camera and Surveyor software (Version 9.0.2.5, Objective Imaging, Kansasville, WI).

### Neurodevelopmental assessments

The neurodevelopmental assessments were administered between approximately 1300 and 1500 hours at seven, 14, 21, 28 (+/− 2) DOL, with the day of birth considered DOL one. Testing occurred in rooms with decreased sensory stimuli to support infant regulation. Throughout the assessment and analysis, the testers were blinded to the infant’s exposure to prevent any bias during collection and analysis.

We evaluated neonatal macaque neurobehavior with a well-validated assessment developed for infant rhesus macaques, termed the Schneider Neonatal Assessment for Primates (SNAP) (37–41), which we have previously used to define neonatal development in prenatally ZIKV-exposed infants (42). The SNAP comprises 29 test items in the neurodevelopmental areas of interest and is made up of the Orientation, Motor maturity and activity, Sensory responsiveness, and State control developmental constructs. Two trained examiners were present for all neurobehavioral testing and scoring to ensure test administration reliability (>95% agreement between examiners). Items were administered in a consistent sequence across all animals to optimize performance and decrease handling time. Assessments were hand-scored during administration and forms were transferred to electronic versions by Qualtrics Survey Software (Qualtrics, Provo, UT). The ratings were based on a five point Likert scale ranging from zero to two. Higher scores reflect optimal scores; variables in which higher numbers do not reflect optimal scores were reverse coded. Test items which represent repetitions of the same skill, such as right, left, up, and down orientation, were averaged together before calculating the average of all the test items within a construct, as described previously (42).

### Locomotion assessments

Immediately following the SNAP assessment, the infant was transported to a new testing room for a quantitative measurement of the animals locomotion using the CatWalk XT version 10.6 (Noldus Information Technology, Wageningen, The Netherlands). Human testers placed the infant into the opening of the CatWalk to walk through a 130 cm long walkway. As the animal walked across and applied pressure on the walkway the green LEDs were refracted on the red background to capture steps. A high-speed digital camera recorded the infant as they moved across the walkway into a nesting box or was picked up by a human tester. The human tester placed the infant back into the opening of the CatWalk until at least three usable runs were collected or the infant was determined to be unable to complete or refused the task. A usable run was defined as having at least two consecutive footfalls per limb on the walkway without stopping. If the infant was unable to walk through, the tester attempted to train the infant by putting them partially through the CatWalk system or placing their blanket in the system as a reinforcer. Data was saved to a USB drive and then immediately transferred to a secure shared research folder.

A group mean and standard deviation for the three runs for each locomotion parameter with Noldus software (Noldus Information Technology, Wageningen, The Netherlands). We evaluated the animals for interlimb coordination and for temporal gait parameters using duty cycle, balance using base of support, and gait maturity categorizing by a diagonal walking pattern. Duty cycle (stand/stand+swing) is the percentage of time the infant had their limb on the walkway (stand) during a step cycle. Base of support includes both the distance between front right and left limb placements and the distance between left and right hind limb placement. The diagonal walking pattern is defined by when a hindlimb paw touches the ground and is closely followed by or occurs simultaneously with the contralateral limb paw (43,44). The speed (distance/time) in which the infant traversed the CatWalk was also measured. All temporal gait parameters were reported as percentages of the step cycle to control for variance in animal velocity, limb length, and body weight.

### Statistical analyses

Overall plasma vRNA loads were calculated for dams receiving a high dose of ZIKV-DAK (1×10^8^ PFU/ml) and historical controls that received a lower dose (1×10^4^ PFU/ml) (24) using the trapezoidal method to calculate the area under the curve (AUC) in R Studio v. 1.4.1717. AUC values were then compared between groups (high-dose versus low-dose) using a t-test. Peak plasma vRNA loads, as well as the duration of positive vRNA detection as defined previously, were also compared between dams receiving either a high dose or low dose of virus inoculum using Wilcoxon rank sum tests with continuity correction using R Studio v. 1.4.1717. The time to peak plasma vRNA load was also compared between dams. Time to peak was analyzed using a linear mixed effects model with the day of peak plasma vRNA load and group (high-dose or low-dose) as the fixed effects and animal ID as the random effect to account for variability by individual within each group using the <u>nlme package in R Studio v. 1.4.1717</u>. The distribution of model residuals was approximately normal. For all analyses of plasma vRNA loads, reported p-values are two-sided and P<0.05 was used to define statistical significance.

In-utero growth data were analyzed by calculating Z-scores which were derived from fitting a spline regression for each outcome (HC, BPD, femur length, abdominal circumference) using the normative reference data described by Tarantal et al. (32). The first available measurement for each outcome was taken as baseline and changes in Z-scores from baseline were calculated for each fetus and analyzed using a linear mixed effects model with animal-specific random effects and an autoregressive correlation structure of order one. Slope parameters for changes in Z-score were calculated for each group (high-dose, low-dose, and mock) and reported with corresponding two-sided 95% confidence intervals (CI) (S10 Fig, S11 Table, S12 Table). Z-scores were not constructed for ratio outcomes (HC-femur length ratio, BPD-femur length ratio) instead, the raw values were log-transformed and analyzed directly.

Quantitative infant variables measured at birth were summarized as means and standard deviations and compared between experimental groups (high dose, low dose, and mock-inoculated) using oneway analysis of variance (ANOVA) with post-hoc pairwise comparisons. Infant gender distribution was compared between experimental groups using Fisher’s exact test. Birth measurements were compared between experimental conditions using analysis of covariance (ANCOVA) with gestational day, dam age, and weight as covariates. Residual and normal probability plots were examined to verify the model assumptions. The results were reported in terms of model-adjusted means along with the corresponding 95% confidence intervals (95%CI) (S13 Table, S14 Table, S15 Table).

Longitudinal changes in infant SNAP parameters were analyzed using a linear mixed effects model with animal specific random effects and an autoregressive correlation structure was used to compare subscale scores between groups each week. Group, week and the interaction effect between group and week were included as factors in the linear mixed effects model. Days before placement with a female and birth weight were included as covariates. Residual plots and histograms were examined to verify the distribution assumptions. All reported P-values are two-sided and P<0.05 was used to define statistical significance. Statistical analyses were conducted using SAS software (SAS Institute, Cary NC), version 9.4 (S19 Fig, S21 Table).

Longitudinal changes in infant gait parameters were analyzed using a linear mixed-effects model with animal-specific random effects and an autoregressive correlation structure of order one to account for the repeated measures from week two to week four. Group, week and the two-way interaction effect between group and week were included as the factors. Number of days before placement with a female, gestational age, and birth weight and speed were included as covariates (speed was excluded when comparing speed between groups). Sex was not included as a covariate because the control group was all female and it was not possible to compare groups with this coviarate. Model assumptions were validated by examining residual plots. The results are reported in terms of adjusted means and the corresponding 95% confidence intervals. Success rates were compared each week between groups using Fisher’s exact test. Diagonal walking pattern comparisons were adjusted by the number of days before placement with a female using an exact, simulation based, logistic regression approach with binomial outcomes (it was not computationally possible to adjust the analysis by birth weight or gestational age). The Wilson score method was used to construct 95% confidence intervals for the success rates (S20 Fig, S22 Table).

### Data availability

All of the data used in this study can also be found at https://github.com/jrrosinski/High-dose-african-lineage-zikv-in-pregnant-rhesus-macaques. In the future, primary data that support this study will also be available at the Zika Open Research Portal (https://openresearch.labkey.com/project/ZEST/begin.view). Data for the high-dose ZIKV-DAK-infected cohort can be found under study ZIKV-046; data for the low-dose ZIKV-DAK infected cohort can be found under study ZIKV-030, ZIKV-PR and mock-inoculated cohorts can be found under ZIKV-044.

---

**S1 Fig. Demographic characteristics of the 17 pregnant rhesus macaques in this study**. Animals were inoculated subcutaneously (SQ) with either 1×10^8^ PFU ZIKV-DAK, 1×10^4^ PFU ZIKV-DAK, or PBS between GD 41 and GD 50.

**S2 Fig. Maternal temperature and weight of dams receiving high-dose (1×10^8^ PFU) ZIKV-DAK**. (A) Maternal temperatures over time. (B) Maternal weights over time. Dams with early fetal loss are marked with an asterisk and were not monitored after cesarean section.

**S3 Fig. Viral loads in plasma samples detected by ZIKV-specific RT-qPCR**. (A) Viral loads through 60 DPI in high-dose dams. Dams with early fetal loss are marked with a single asterisk. (B) Viral loads through 60 DPI in low-dose dams.

**S4 Fig. Replication of ZIKV-DAK in (A) urine and (B) saliva of dams receiving 1×10^8^ PFU ZIKV-DAK**. Viral loads were measured by ZIKV-specific RT-qPCR.

**S5. Replication kinetics of ZIKV-DAK in high-dose dams, grouped by birth outcome**. Viral loads were measured in plasma samples by ZIKV-specific qRT-PCR. (A) Comparison of area under the curve (AUC). For all graphs in parts B-E, the mean value is shown with error bars representing standard deviation. (B) Days post-infection of the last positive plasma vRNA load. (C) Peak plasma viral load in copies/ml plasma. (D) Day post-infection of peak plasma viral load. *** Represents a p-value of <0.001, while no asterisk represents no statistical difference.

**S6 Fig. All animals developed robust neutralizing antibody titers after inoculation**. Plaque reduction neutralization tests were performed on serum samples collected zero days post infection (DPI) and between 21 and 28 DPI from animals infected with high-dose (1×10^8^ PFU) or a low dose (10^4^ PFU) of ZIKV-DAK. Dams with early fetal loss are marked with an asterisk.

**S7 Fig. Maternal serum IgM titers measured by ELISA in high-dose animals**. Levels of IgM antibodies were measured in maternal serum samples at zero, seven, 14, 17 and 21 or 24 DPI. Values are normalized to AbCam units for comparison based on the manufacturer’s recommendation (see materials and methods).

**S8 Table. In utero imaging observations and interpretations from dams that received high-dose ZIKV-DAK.**

**S9 Fig. Fetal heart rate throughout gestation**. Fetal heart rate was monitored biweekly by ultrasound on high-dose (1×10^8^ PFU ZIKV-DAK) dams to confirm fetal viability. The horizontal, dotted lines on the graph represent the range of normal heart rates for fetuses at WNPRC. Cases of early fetal loss are marked with an asterisk.

**S10 Fig. In-utero fetal growth from weekly sonographic imaging**. Normative data generated by Tarantal at CNPRC were used to calculate Z-scores for each animal. Open circles represent the change in Z-score from baseline for each animal. Solid lines show the growth trajectories for each group and were quantified by calculating regression slope parameters from baseline using a linear mixed-effects model with animal-specific random effects and an autoregressive correlation structure.

**S11 Table. Slope parameters for changes in z-scores for in-utero measurements across gestation, stratified by group**. Z-scores are log-transformed ratios. Animals in high-dose ZIKV-DAK, low-dose ZIKV-DAK, and mock groups were compared.

**S12 Table. Pairwise comparisons between groups of slopes for z-scores of in-utero measurements**. Animals in high-dose ZIKV-DAK, low-dose ZIKV-DAK, and mock groups were compared.

**S13 Table. Statistical analyses comparing weights, head circumference (HC), biparietal diameter (BPD) and weight**. Analysis was performed between infants in high-dose ZIKV-DAK and mock groups. Because gender was confounded with group it was not included as a covariate in this analysis.

**S14 Table. Statistical analyses comparing APGAR scores at one, five, and 10 minutes of life**. Comparisons of APGAR scores were made between infants in high-dose, low-dose, and mock groups. Because gender was confounded with group it was not included as a covariate in this analysis.

**S15 Table. Statistical analysis comparing at-birth demographic characteristics**. Characteristics of gestational day (GD), dam weight, dam age, and fetal gender were compared between mock (n=4), high-dose (n=5), and low-dose (n=4) groups.

**S16 Fig. Transmural infarcts in the placentas of dams with early fetal loss**. (A) Placental infarction in 046-103 and (B) 046-107. Placental tissues were stained with H&E and are imaged at 4X magnification. Black boxes denote areas of infarct.

**S17 Table. Morphological diagnoses of MFI and fetal tissue from animals in high-dose, low-dose, and control groups.**

**S18 Fig. Detection of ZIKV RNA by in situ hybridization in fetal body tissues**. Representative images of ZIKV RNA distribution in fetal tissues from the three cases of early fetal loss (excluding the head). Foci of ZIKV RNA were detected in the neuropil of (A) 046-503 (boxed), (B) 046-504 (arrows), and (C) 046-507; the spinal cord of (D) 046-503 (arrows), (E) 046-504 (boxed), and (F) 046-507 (arrows); the intestines of (G) 046-503 (boxed, arrows) and (I) 046-507 (boxed, arrows); and the periosteum of (J) 046-503 (boxed, arrows) and (K) 046-507 (arrows). ZIKV RNA is shown in red.

**S19 Fig. Neonatal neurodevelopment measured by SNAP in the first month of life**. Neurodevelopment was measured by SNAP at seven, 14, 21, and 28 days of life for high-dose ZIKV-DAK exposed (ZIKV-HD) and control infants. Scores in the (A) Orientation, (B) Motor Maturity and Activity, (C) Sensory Responsiveness, and (D) State control constructs are illustrated while controlling for birth weight and days reared in nursery conditions. Animals were rated on a five point Likert scale ranging from zero to two with higher scores reflecting optimal scores. For all graphs shown the results are reported in terms of model-adjusted means along with the corresponding 95% confidence intervals (95% CI).

**S20 Fig. Locomotion in the first month of life measured by the CatWalk**. (A) Screenshot of the infant completing a run in the CatWalk Noldus XT measuring footfalls on a pressure plate that are labeled showing bilateral front (RF, LF) and hind (RH, LH) limbs. For all graphs shown in C-F, the results are reported in terms of model-adjusted means along with the corresponding 95% confidence intervals (95% CI). (B) Visual representation of gait variables including duty cycle (stand/stand + swing), base of support, and speed (cm/s), (C) % of runs in mature locomotion pattern, (D) Average speed across runs, (E) Base of support defined between the left and right foot/hand prints, and F) Duty cycle defined as the % time spent on each limb.

**S21 Table. Comparison of neonatal development with the Schneider Neonatal Assessment Protocol (SNAP)**. Infants exposed to high-dose ZIKV-DAK are compared to control infants.

**S22 Table. Comparison of gait development with the Nodulus CatWalk**. Infants exposed to high-dose ZIKV-DAK are compared to control infants.

## Bibliography

1. Duffy MR, Chen T-H, Hancock WT, Powers AM, Kool JL, Lanciotti RS, et al. Zika virus outbreak on Yap Island, Federated States of Micronesia. N Engl J Med. 2009 Jun 11;360(24):2536–43.

2. Lanciotti RS, Lambert AJ, Holodniy M, Saavedra S, Signor LDCC. Phylogeny of Zika Virus in Western Hemisphere, 2015. Emerg Infect Dis. 2016 May;22(5):933–5.

3. Zhang Q, Sun K, Chinazzi M, Pastore Y Piontti A, Dean NE, Rojas DP, et al. Spread of Zika virus in the Americas. Proc Natl Acad Sci U S A. 2017 May 30;114(22):E4334–43.

4. Passemard S, Kaindl AM, Verloes A. Microcephaly. Handb Clin Neurol. 2013;111:129–41.

5. Rasmussen SA, Jamieson DJ, Honein MA, Petersen LR. Zika Virus and Birth Defects — Reviewing the Evidence for Causality [Internet]. Vol. 374, New England Journal of Medicine. 2016. p. 1981–7. Available from: http://dx.doi.org/10.1056/nejmsr1604338

6. Schuler-Faccini L, Ribeiro EM, Feitosa IML, Horovitz DDG, Cavalcanti DP, Pessoa A, et al. Possible Association Between Zika Virus Infection and Microcephaly - Brazil, 2015. MMWR Morb Mortal Wkly Rep. 2016 Jan 29;65(3):59–62.

7. Victora CG, Schuler-Faccini L, Matijasevich A, Ribeiro E, Pessoa A, Barros FC. Microcephaly in Brazil: how to interpret reported numbers? Lancet. 2016 Feb 13;387(10019):621–4.

8. Moore CA, Staples JE, Dobyns WB, Pessoa A, Ventura CV, Fonseca EB da, et al. Characterizing the Pattern of Anomalies in Congenital Zika Syndrome for Pediatric Clinicians. JAMA Pediatr. 2017 Mar 1;171(3):288–95.

9. Esser-Nobis K, Aarreberg LD, Roby JA, Fairgrieve MR, Green R, Gale M Jr. Comparative Analysis of African and Asian Lineage-Derived Zika Virus Strains Reveals Differences in Activation of and Sensitivity to Antiviral Innate Immunity. J Virol [Internet]. 2019 Jul 1;93(13). Available from: http://dx.doi.org/10.1128/JVI.00640-19

10. Haddow AD, Schuh AJ, Yasuda CY, Kasper MR, Heang V, Huy R, et al. Genetic characterization of Zika virus strains: geographic expansion of the Asian lineage. PLoS Negl Trop Dis. 2012 Feb 28;6(2):e1477.

11. Faye O, Freire CCM, Iamarino A, Faye O, de Oliveira JVC, Diallo M, et al. Molecular evolution of Zika virus during its emergence in the 20(th) century. PLoS Negl Trop Dis. 2014 Jan 9;8(1):e2636.

12. Beaver JT, Lelutiu N, Habib R, Skountzou I. Evolution of Two Major Zika Virus Lineages: Implications for Pathology, Immune Response, and Vaccine Development. Front Immunol. 2018 Jul 18;9:1640.

13. Dudley DM, Van Rompay KK, Coffey LL, Ardeshir A, Keesler RI, Bliss-Moreau E, et al. Miscarriage and stillbirth following maternal Zika virus infection in nonhuman primates. Nat Med. 2018 Aug;24(8):1104–7.

14. Cauvin AJ, Peters C, Brennan F. Chapter 19 - Advantages and Limitations of Commonly Used Nonhuman Primate Species in Research and Development of Biopharmaceuticals. In: Bluemel J, Korte S, Schenck E, Weinbauer GF, editors. The Nonhuman Primate in Nonclinical Drug Development and Safety Assessment. San Diego: Academic Press; 2015. p. 379–95.

15. Jaeger AS, Murrieta RA, Goren LR, Crooks CM, Moriarty RV, Weiler AM, et al. Zika viruses of African and Asian lineages cause fetal harm in a mouse model of vertical transmission. PLoS Negl Trop Dis. 2019 Apr;13(4):e0007343.

16. Duggal NK, Ritter JM, McDonald EM, Romo H, Guirakhoo F, Davis BS, et al. Differential neurovirulence of African and Asian genotype Zika virus isolates in outbred immunocompetent mice. Am J Trop Med Hyg. 2017 Nov;97(5):1410–7.

17. Aubry F, Jacobs S, Darmuzey M, Lequime S, Delang L, Fontaine A, et al. Recent African strains of Zika virus display higher transmissibility and fetal pathogenicity than Asian strains. Nat Commun. 2021 Feb 10;12(1):916.

18. Sheridan MA, Balaraman V, Schust DJ, Ezashi T, Roberts RM, Franz AWE. African and Asian strains of Zika virus differ in their ability to infect and lyse primitive human placental trophoblast. PLoS One. 2018 Jul 9;13(7):e0200086.

19. Udenze D, Trus I, Berube N, Gerdts V, Karniychuk U. The African strain of Zika virus causes more severe in utero infection than Asian strain in a porcine fetal transmission model. Emerg Microbes Infect. 2019;8(1):1098–107.

20. Crooks CM, Weiler AM, Rybarczyk SL, Bliss M, Jaeger AS, Murphy ME, et al. African-lineage Zika virus replication dynamics and maternal-fetal interface infection in pregnant rhesus macaques [Internet]. bioRxiv. 2020 [cited 2021 Jun 10]. p. 2020.11.30.405670. Available from: https://www.biorxiv.org/content/10.1101/2020.11.30.405670v1

21. Adams Waldorf KM, Stencel-Baerenwald JE, Kapur RP, Studholme C, Boldenow E, Vornhagen J, et al. Fetal brain lesions after subcutaneous inoculation of Zika virus in a pregnant nonhuman primate. Nat Med. 2016 Nov;22(11):1256–9.

22. Styer LM, Kent KA, Albright RG, Bennett CJ, Kramer LD, Bernard KA. Mosquitoes inoculate high doses of West Nile virus as they probe and feed on live hosts. PLoS Pathog. 2007 Sep 14;3(9):1262–70.

23. Gubler DJ, Rosen L. A simple technique for demonstrating transmission of dengue virus by mosquitoes without the use of vertebrate hosts. Am J Trop Med Hyg. 1976 Jan;25(1):146–50.

24. Crooks CM, Weiler AM, Rybarczyk SL, Bliss M, Jaeger AS, Murphy ME, et al. African-Lineage Zika Virus Replication Dynamics and Maternal-Fetal Interface Infection in Pregnant Rhesus Macaques. J Virol. 2021 Jul 26;95(16):e0222020.

25. Nutt C, Adams P. Zika in Africa—the invisible epidemic? Lancet. 2017 Apr 22;389(10079):1595–6.

26. Dudley DM, Newman CM, Lalli J, Stewart LM, Koenig MR, Weiler AM, et al. Infection via mosquito bite alters Zika virus tissue tropism and replication kinetics in rhesus macaques [Internet]. Vol. 8, Nature Communications. 2017. Available from: http://dx.doi.org/10.1038/s41467-017-02222-8

27. Pomar L, Lambert V, Matheus S, Pomar C, Hcini N, Carles G, et al. Prolonged maternal Zika viremia as a marker of adverse perinatal outcomes. Emerg Infect Dis. 2021 Feb;27(2):490–8.

28. Aragao MFVV, Brainer-Lima AM, Holanda AC, van der Linden V, Vasco Aragão L, Silva Júnior MLM, et al. Spectrum of Spinal Cord, Spinal Root, and Brain MRI Abnormalities in Congenital Zika Syndrome with and without Arthrogryposis. AJNR Am J Neuroradiol. 2017 May;38(5):1045–53.

29. Ramalho FS, Yamamoto AY, da Silva LL, Figueiredo LTM, Rocha LB, Neder L, et al. Congenital Zika Virus Infection Induces Severe Spinal Cord Injury. Clin Infect Dis. 2017 Aug 15;65(4):687–90.

30. de Fatima Vasco Aragao M, van der Linden V, Brainer-Lima AM, Coeli RR, Rocha MA, Sobral da Silva P, et al. Clinical features and neuroimaging (CT and MRI) findings in presumed Zika virus related congenital infection and microcephaly: retrospective case series study. BMJ. 2016 Apr 13;353:i1901.

31. National Research Council, Division on Earth and Life Studies, Institute for Laboratory Animal Research, Committee for the Update of the Guide for the Care and Use of Laboratory Animals. Guide for the Care and Use of Laboratory Animals: Eighth Edition. National Academies Press; 2011. 246 p.

32. Tarantal AF. Ultrasound Imaging in Rhesus (Macaca mulatta) and Long-tailed (Macaca fascicularis) Macaques: Reproductive and Research Applications [Internet]. The Laboratory Primate. 2005. p. 317–52. Available from: http://dx.doi.org/10.1016/b978-012080261-6/50020-9

33. Hansen SG, Piatak M Jr, Ventura AB, Hughes CM, Gilbride RM, Ford JC, et al. Immune clearance of highly pathogenic SIV infection. Nature. 2013 Oct 3;502(7469):100–4.

34. Lanciotti RS, Kosoy OL, Laven JJ, Velez JO, Lambert AJ, Johnson AJ, et al. Genetic and serologic properties of Zika virus associated with an epidemic, Yap State, Micronesia, 2007. Emerg Infect Dis. 2008 Aug;14(8):1232–9.

35. Breitbach ME, Newman CM, Dudley DM, Stewart LM, Aliota MT, Koenig MR, et al. Primary infection with dengue or Zika virus does not affect the severity of heterologous secondary infection in macaques. PLoS Pathog. 2019 Aug;15(8):e1007766.

36. Lindsey HS, Calisher CH, Mathews JH. Serum dilution neutralization test for California group virus identification and serology. J Clin Microbiol. 1976 Dec;4(6):503–10.

37. Laughlin NK, Lasky RE, Giles NL, Luck ML. Lead effects on neurobehavioral development in the neonatal rhesus monkey (Macaca mulatta). Neurotoxicol Teratol. 1999 Nov;21(6):627–38.

38. Schneider ML, Moore CF, Kraemer GW, Roberts AD, DeJesus OT. The impact of prenatal stress, fetal alcohol exposure, or both on development: perspectives from a primate model. Psychoneuroendocrinology. 2002 Jan;27(1-2):285–98.

39. Schneider ML, Roughton EC, Koehler AJ, Lubach GR. Growth and development following prenatal stress exposure in primates: an examination of ontogenetic vulnerability. Child Dev. 1999 Mar;70(2):263–74.

40. Levin ED, Schneider ML, Ferguson SA, Schantz SL, Bowman RE. Behavioral effects of developmental lead exposure in rhesus monkeys. Dev Psychobiol. 1988 May;21(4):371–82.

41. Coe CL, Lubach GR, Crispen HR, Shirtcliff EA, Schneider ML. Challenges to maternal wellbeing during pregnancy impact temperament, attention, and neuromotor responses in the infant rhesus monkey. Dev Psychobiol. 2010 Nov;52(7):625–37.

42. Koenig MR, Razo E, Mitzey A, Newman CM, Dudley DM, Breitbach ME, et al. Quantitative definition of neurobehavior, vision, hearing and brain volumes in macaques congenitally exposed to Zika virus. PLoS One. 2020 Oct 22;15(10):e0235877.

43. Dunbar DC. Locomotor behavior of rhesus macaques (Macaca mulatta) on Cayo Santiago. P R Health Sci J. 1989 Apr;8(1):79–85.

44. Shapiro LJ, Cole WG, Young JW, Raichlen DA, Robinson SR, Adolph KE. Human quadrupeds, primate quadrupedalism, and Uner Tan Syndrome. PLoS One. 2014 Jul 16;9(7):e101758.

